# Machine learning elucidates electrophysiological properties predictive of multi- and single-firing human and mouse dorsal root ganglia neurons

**DOI:** 10.1101/2024.06.03.597213

**Authors:** Nesia A. Zurek, Sherwin Thiyagarajan, Reza Ehsanian, Aleyah E. Goins, Sachin Goyal, Mark Shilling, Christophe G. Lambert, Karin N. Westlund, Sascha R.A. Alles

## Abstract

Human and mouse dorsal root ganglia (hDRG and mDRG) neurons are important tools in understanding the molecular and electrophysiological mechanisms that underlie nociception and drive pain behaviors. One of the simplest differences in firing phenotypes is that neurons are single-firing (exhibit only one action potential) or multi-firing (exhibit 2 or more action potentials). To determine if single- and multi-firing hDRG exhibit differences in intrinsic properties, firing phenotypes, and AP waveform properties, and if these properties could be used to predict multi-firing, we measured 22 electrophysiological properties by whole-cell patch-clamp electrophysiology of 94 hDRG neurons from 6 male and 4 female donors. We then analyzed the data using several machine learning models to determine if these properties could be used to predict multi-firing. We used 1000 iterations of Monte Carlo Cross Validation to split the data into different train and test sets and tested the Logistic Regression, k-Nearest Neighbors, Random Forest, Supported Vector Classification, and XGBoost machine learning models. All models tested had a greater than 80% accuracy on average, with Supported Vector Classification and XGBoost performing the best. We found that several properties correlated with multi-firing hDRG neurons and together could be used to predict multi-firing neurons in hDRG including a long decay time, a low rheobase, and long first spike latency. We also found that the hDRG models were able to predict multi-firing with 90% accuracy in mDRG. Targeting the neuronal properties that lead to multi-firing could elucidate better targets for treatment of chronic pain.

## Introduction

Neuronal hyperexcitability is a hallmark of chronic pain and understanding the electrophysiological mechanisms that lead to neuronal excitability is crucial for the development of pain treatments (Berta et al., 2017; Alles and Smith, 2018). Sensory neurons isolated from human and mouse dorsal ganglia (hDRG-N and mDRG-N) are important tools in the study of neuronal excitability, chronic pain, and nociception (Davidson et al., 2014; Zhang et al., 2017; Emery and Ernfors, 2018; North et al., 2019; Zheng et al., 2019; Zurek et al., 2024). While the transcriptomes of hDRG and mDRG have been compared (Ray et al., 2018), few studies have compared the electrophysiological features of these tools. Single-firing DRG-N that fire only one action potential (AP) are considered less excitable than multi-firing DRG-N that fire 2 or more APs. Using patch-clamp electrophysiology, we measured neuronal excitability of hDRG-N and mDRG-N in current clamp mode. We then compared electrophysiological properties that determine multi- or single-firing in DRG-N using sophisticated computational approaches. The rationale for this methodology is to build upon understanding of electrophysiological properties of sensory neurons with machine learning tools.

Machine learning is becoming a popular tool in data analysis, tool refinement, and prediction in understanding molecular, electrophysiological, and physiological mechanisms in pain research (North et al., 2019; Gonzalez et al., 2021; Koos et al., 2021; Ingram et al., 2023; Nagaraja et al., 2023). In this study, we aimed to understand if there are electrophysiological differences between single- and multi-firing hDRG-N and mDRG-N, and if these differences are predictive of neuronal excitability. Factors governing excitability might thereby be targets for modulation with therapies. Additionally, these improved tools can lead to automated, and therefore more efficient, ways to evaluate future electrophysiological data. The benefits of automating data analysis are better data transparency and reproducibility, while minimizing the need for obtaining additional live tissue.

In this study, we compared the electrophysiological features of multi- and single-firing hDRG-N and mDRG-N and applied several machine learning algorithms to elucidate which combinations of features were most predictive of multi-firing neurons. Because we have collected a relatively high number of hDRG-N recordings from a diverse demographic of donors (Zurek et al., 2024), we aimed to see if machine learning algorithms can predict whether an hDRG-N will be single- or multi-firing based on other intrinsic, phenotypic, and AP waveform electrophysiological features. While many machine learning studies use a single model to make predictions, we used several different models. We used Monte Carlo Cross Validation (MCCV) simulations to iterate the train and test data split to obtain average model accuracies (Shan, 2022) and extract the most important electrophysiological features. Finally, we aimed to see if the machine learning models that we generated were also able to predict multi-firing mDRG-N so as to compare species differences. While we found several electrophysiology features were correlated with multi-firing cells, the machine learning models converged on just a few of those features as being the most predictive.

This study will improve understanding of the electrophysiological mechanisms of DRG neurons. Our machine learning algorithms paradoxically show few species differences between mouse and human DRG neuron electrophysiology under baseline conditions. These are important findings for the study of neuronal excitability in the context of pain therapeutic development.

## Materials and Methods

### hDRG-N Culture

hDRG-N culture was performed as previously described (Valtcheva et al., 2016; Zurek et al., 2024). hDRG were obtained from consenting recently deceased organ donors at University of New Mexico Hospital in coordination with New Mexico Donor Services. Study activities were approved by the Human Research Review Committee at the University of New Mexico Health Sciences Center; approval numbers #21-412 or #23-205. Cultures of hDRG-N were prepared as described previously and cultured for up to 11 days *in vitro* (DIV) (Zurek et al., 2024) and electrophysiological recordings took place between DIV 3 to 11. We used high-quality recordings where all 22 electrophysiological properties could be extracted from 94 untreated hDRG-N. The data from 8 of 10 donors was previously published in (Zurek et al., 2024). Donor demographics are described in Table 1.

**Table 1:** Donor demographics. Human DRG tissue was obtained from ethically consented, recently deceased donors. This analysis used tissues from 4 female and 6 male donors aged 21 to 67 years.

| **hDRG Donors** | **Sex** | **Age (Years)** | **BMI** | **Race** | **Number of Recordings** |
| --- | --- | --- | --- | --- | --- |
| 1 | F | 22 | 22 | White | 1 |
| 2 | F | 51 | 30 | White | 13 |
| 3 | M | 23 | 26 | Hispanic | 14 |
| 4 | M | 67 | 34 | Native | 7 |
| 5 | M | 55 | 19 | White | 2 |
| 6 | M | 27 | 24 | Hispanic | 2 |
| 7 | M | 21 | 22 | White | 13 |
| 8 | F | 37 | 18 | Hispanic | 18 |
| 9 | M | 51 | 27 | Unknown | 3 |
| 10 | F | 23 | 27 | Hispanic | 21 |
| **94 Total Recordings from 10 donors** |  |  |  |  |  |

### Mouse DRG cultures

All animal procedures were compliant with the NIH Guide for the Care and Use of Laboratory Animals and followed ARRIVE guidelines. Studies are approved by the Institutional Care and Use Committee of the University of New Mexico Health Sciences Center (IACUC #23-201364-HSC, 5-13-2024). Mouse DRG cultures were performed as previously described with some modifications (Kunamneni et al., 2023). Briefly, lumbar DRG were collected and put into an enzymatic solution containing sterile, magnesium/calcium-free HBSS, papain, dispase, and collagenase. The enzymatic digestion was carried out at 37°C, 5% CO2 for 40 minutes with trituration every 20 minutes. Complete DMEM-based media (2mL DMEM supplemented with 10% fetal bovine serum, and 1% antibacterial/antimycotic with 100 units/mL of penicillin, 100 µg/mL of streptomycin, 25 µg/mL of amphotericin B) was added to the enzymatic digestion. The solution was then strained through a 100µm cell strainer and rinsed several times with an additional 6mL complete media. The digested DRG cell suspension was rinsed by gentle centrifugation at 300Xg for 5 minutes and resuspended in 1mL complete media. 125µL of the mouse DRG cell suspension was added to each 12mm coverslip pre-coated with poly-d-lysine (Neuvitro, Camas, WA) and coated with additional 50µg/mL laminin, allowed to attach for 30-60 minutes before gently flooding the wells with enough media to fill each well, 1-2mL for a 12 well plate. Electrophysiological recordings were done 18-24 hours after mDRG culture completion.

### Whole-cell Patch-clamp Electrophysiology

Whole-cell patch-clamp electrophysiology was performed as previously described (Zurek et al., 2024). Recordings were done at room temperature, with the recording chamber perfused with artificial cerebrospinal fluid (aCSF) (Zurek et al., 2024). Neurons were identified with differential interference contrast optics connected to an IR-2000 digital camera (Dage MTI, Indiana City, MI) or an Olympus digital camera. Cell diameter was measured using Dage MTI camera software or ImageJ (NIH, Bethesda, MD). Current-clamp recordings were performed using a Multiclamp 700B (Molecular Devices, San Jose, CA). Signals were acquired as previously described using a Digidata 1550B converter (Molecular Devices, San Jose, CA), and recorded using Clampex 11 software (Molecular Devices, San Jose, CA). Patch pipettes with electrode resistance of 3-7 MΩ were made fresh with a Zeitz puller (Werner Zeitz, Martinsried, Germany) from borosilicate thick glass (GC150F, Sutter Instruments, Novato, CA). Intracellular patch-pipette solution was the same as previously described (Zurek et al., 2024). Cells that did not fire APs or had an RMP of >-35mV were excluded from further analysis. Bridge balance was applied for all recordings.

Analysis was performed in Easy Electrophysiology v.2.5.1 (London, UK), Clampfit 11.2 (Molecular Devices, San Jose, CA), and the Python v3.12 package pyABF (Harden, 2022). All statistical analysis was performed using GraphPad Prism v10.0.2 (Boston, MA). Error bars denote mean ± standard error of the mean (SEM) unless otherwise specified.

### Electrophysiology Intrinsic Properties Analysis Methods

Electrophysiological analysis was done as previously described and used recordings from 8 human donors published in (Zurek et al., 2024) and data from 2 additional donors not previously published. Current-clamp recordings used a 500 ms current pulse increasing in 10pA increments from −100 pA in increments until they reached inactivation up to 4nA. This recording was used to calculate resting membrane potential (RMP), input resistance (R_in_), sag ratio, rebound firing, rheobase, and first spike latency (FSL) using Easy Electrophysiology Software and pyABF (Harden, 2022; Zurek et al., 2024). Cell capacitance was calculated using the whole-cell capacitance compensation circuit in Multiclamp 700B (Molecular Devices). Normalized rheobase was calculated by dividing rheobase by capacitance. Delayed firing was classified as a neuron that had an FSL >100ms. Presence of visible sag was identified by visual inspection. hDRG-N that fired more than one AP during any current injection step were labeled as multi-firing (65/94 for hDRG-N, 14/20 for mDRG-N) while neurons that fired only one AP were labeled single-firing (29/94 for hDRG-N, 6/20 for mDRG-N) in our classification system. Spontaneous activity was assessed in current clamp mode with either no current injection or injecting enough current injection to hold the membrane voltage at -45mV during a 30s recording. If the neuron had at least one AP during the 30s it was recorded as having spontaneous activity. AP waveform properties were calculated using Easy Electrophysiology as previously described (Zurek et al., 2024). Only DRG-N which we were able to accurately measure all the properties were considered for further analysis. The same analysis techniques were used both hDRG-N and mDRG-N.

Statistics comparing all electrophysiological properties between single- and multi-firing hDRG-N were performed in GraphPad Prism v10.0.2 using a Mann-Whitney test or a Fisher’s Exact test.

### Pearson Correlation Matrices

Pearson Correlation Matrices for both hDRG-N and mDRG-N were generated in Python v3.12 using the Seaborn package for all electrophysiological properties described (Waskom, 2021). Jupyter notebooks was used to write and share code and PANDAS was used for data processing (McKinney, 2010; Kluyver et al., 2016).

### Machine Learning Methods

All machine learning methods were performed in Python v3.12 using scikit-learn (Pedregosa et al., 2011), an open-source machine learning package. Jupyter Notebooks was used as a framework for writing, editing, and sharing code across unscaled, scaled, and feature-selected datasets (Kluyver et al., 2016). The open-source package, PANDAS (McKinney, 2010), was used for data processing, scaling, normalization, and analysis. Matplotlib and Seaborn data visualization libraries were used to produce figures (Hunter, 2007; Waskom, 2021). The dataset preprocessing included standardizing the dataset using the standard scaler in scikit-learn, converting categorical variables to binary variables, and feature selection. The unprocessed and preprocessed dataset was then used to tune the hyperparameters of the four models we used: logistic regression (LR), k-nearest neighbors (KNN), random forest (RF), support vector classifier (SVC), and eXtreme Gradient Boost (XGBoost). The tuning was accomplished using GridSearchCV with a five-fold split, a scikit-learn method that performs an exhaustive search over the given hyperparameters to determine which combination of hyperparameters generates the highest accuracy for a given dataset and model. The optimal hyperparameters were then used during the MCCV. Afterwards, the dataset was inputted into MCCV for each of the hyper-parameterized models. The MCCV split the original dataset into a training and testing dataset without replacement where the training dataset contains 80% of the samples and the testing dataset consists of 20% of the samples. Then, the training dataset was used to train the model, which was then applied to the testing dataset.

The predictions that the trained model creates based on the testing dataset are compared with the presence of multi-firing for those cells to produce an accuracy value. The creation of the subsets, training, and testing is repeated 1000 times, allowing us to extract various metrics and attributes of the model across the iterations, resulting in comprehensive measures of the model that account for variations in the training and testing dataset induced by the random selection of the datasets. Utilizing the MCCV, we were able to visualize the distribution of the accuracy, precision, recall and F1 values for each of the models and obtain the corresponding median, mean, standard error of the mean, and range. Additionally, by storing the coefficient or feature importance values for each of the MCCV iterations, we derived the mean weight given to each feature coefficient throughout the iterations. We also used Shapley Additive Explanations (Lundberg and Lee, 2017) to extract feature importance across all models on the 80%/20% split using MCCV. The SHAP values were plotted on beeswarm plots and then bar plots showing the absolute average SHAP values.

### Principal Component Analysis (PCA)

As an alternate approach to understanding feature importance, we used the selected features of hDRG-N to perform a PCA in Python v3.12 using scikit-learn (Pedregosa et al., 2011). Jupyter notebooks were used to write and share code and PANDAS was used for data processing and normalization (McKinney, 2010; Kluyver et al., 2016). We first normalized the data using the ‘normalize’ function, then applied 3 component analysis. We extracted the explained variance and feature importance using scikit-learn and plotted the data using matplot-lib (Hunter, 2007).

## Results

### Intrinsic Properties in multi-versus single-firing hDRG-N

Using patch-clamp electrophysiology in current clamp mode, we compared a variety of intrinsic and phenotypic properties of single- and multi-firing hDRG-N. Single-firing hDRG-N made up 30.8% (29/94) of our data set. Cell diameter measured during clamp recording with an IR-2000 Dage MTI camera (Figure 1A) showed that multi-firing hDRG-N are smaller than single-firing hDRG-N (45.03 ± 0.95µm and 49.69 ± 2.15µm respectively, p=0.0377). Cell capacitance was also significantly different, single-firing cells had an average capacitance 132.3 ± 13.34pF, and multi-firing cells had an average capacitance of 84.58 ± 6.26pF, p=0.0008 (Figure 1B). There was no significant difference in RMP between single and multi-firing cells (Figure 1C). Single-firing cells had a significantly lower input resistance (R_in_), 89.23 ± 13.55MΩ, compared to 311.2 ± 40.25MΩ for multi-firing cells, p<0.0001 (Figure 1D). Rheobase for single firing cells (927.2 ± 103.8pA) was significantly higher than rheobase for multi-firing cells (340.6 ± 58.86pA), p<0.0001 (Figure 1E), though when rheobase was normalized to cell capacitance, there was no significant difference in normalized rheobase (Figure 1F). FSL was sooner in single-firing cells than multi-firing cells (40.64 ± 18.63ms and 141.50 ± 20.26ms respectively, p<0.0001) (Figure 1G). We defined cells with delayed firing as cells with an FSL >100ms and multi-firing cells (37%) were much more likely to exhibit delayed firing than single-firing cells (7%) consistent with the FSL data, p<0.0001 (Figure 1H). There was no significant difference in sag percentage (Figure 1I), however multi-firing cells were more likely, 66%, to show visible sag than single-firing cells, 34%, p<0.0001 (Figure 1J) on visual inspection. There was no significant difference in rebound firing (Figure 1K). We also looked at spontaneous activity of the cells both at rest and when applying enough current to hold the cell at -45mV membrane potential. At rest, only multi-firing cells exhibited any spontaneous activity, 12%, compared to 0% for single firing cells (Figure 1L). Multi-firing cells also had more spontaneous activity when held at -45mV, 55% of multi-firing cells compared to only 7% of single-firing cells (Figure 1M).

**Figure 1:**
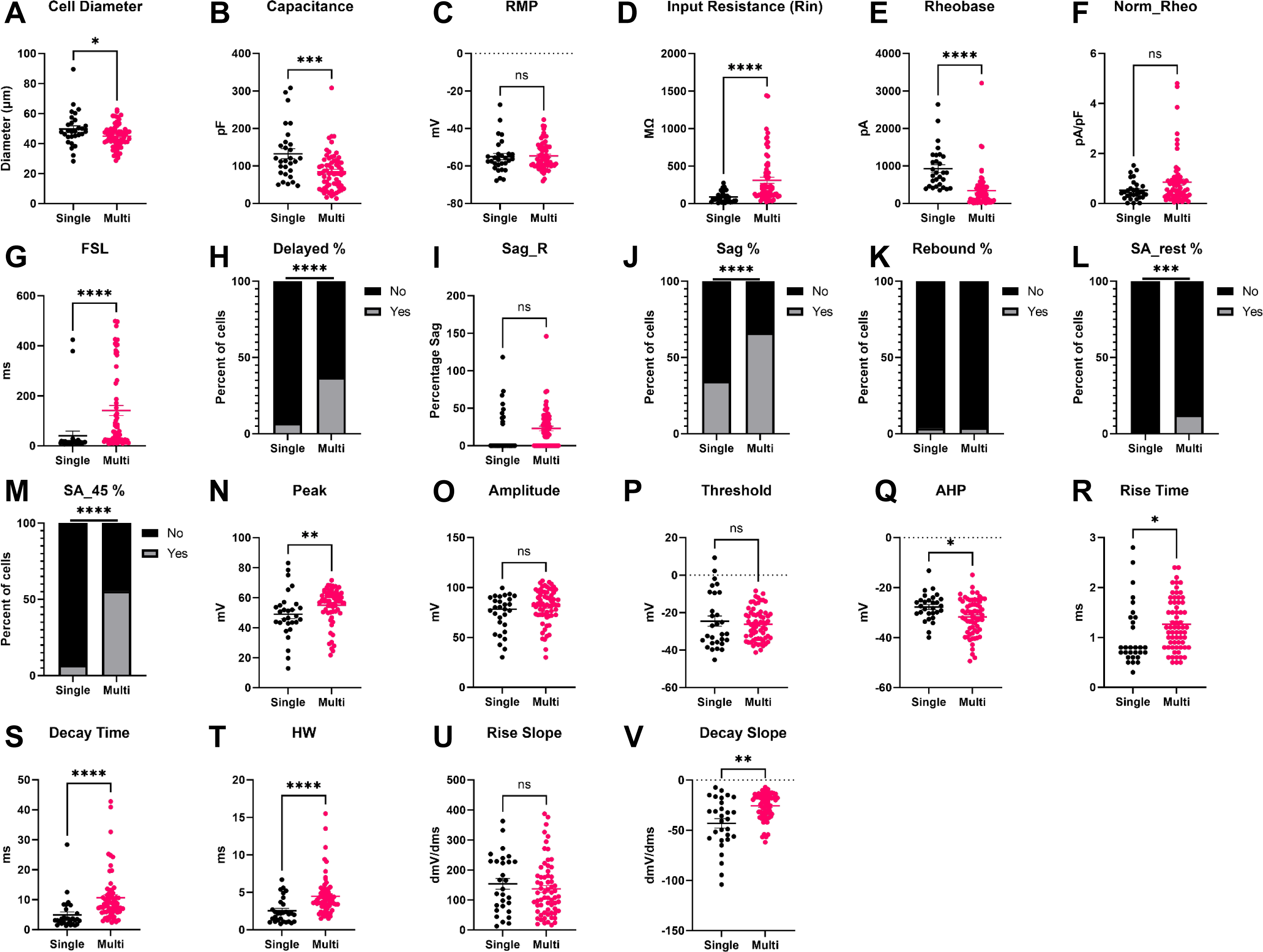
Intrinsic Properties and Firing Phenotypes of hDRG-N neurons in single versus multi-firing cells. Intrinsic properties and firing phenotypes were compared between single and multi-firing cells. A. Cell Diameter, B. Capacitance, C. Resting Membrane Potential (RMP), D. Input Resistance (Rin), E. Rheobase, F. Normalized Rheobase, G. First-Spike Latency (FSL), H. Percent delayed firing, FSL > 100ms, I. Sag Percentage at -100pA current input (Sag_R), J. Percentage cells with visible sag, K. Percentage of cells that have rebound firing. L. Percentage cells with spontaneous activity (SA) at rest, M. Percentage cells with spontaneous activity when held at -45mV. N. AP Peak. O. AP Amplitude. P AP Threshold. Q. AP After hyper-polarization (AHP). R. AP Rise Time, S. AP Decay Time, T. AP half-width. U. Max AP Rise Slope. V. Max AP Decay Slope. *p<0.05, **p<0.01, ***p<0.001, ****p<0.0001 by Mann-Whitney for A, B, C, D, E, F, G, I, N, O, P, Q, R, S, T, U, and V. By Fisher’s Exact test for H, J, K, L, and M .

### Action potential (AP) waveform properties in multi-versus single-firing hDRG-N

#### AP Amplitude

Single- and multi-firing hDRG-N AP waveform properties were analyzed using whole cell patch-clamp electrophysiology in current clamp mode of the rheobase spike. There were no significant differences found in AP amplitude or AP threshold (Figure 1O and 1P). However, there was a significant difference in AP peak. Single-firing hDRG-N had a smaller AP peak of 48.95 ± 2.93mV and multi-firing hDRG-N had an AP peak of 54.95 ± 1.46mV, p=0.0068 (Figure 1N). There was also a difference in AP after hyperpolarization (AHP). Single-firing cells had an AHP of -27.77 ± 1.00mV and multi-firing cells had and AHP of -31.79 ± 0.90mV, p=0.0219 (Figure 1Q).

#### AP Duration

AP duration of the rheobase in hDRG-N was analyzed and multi-firing cells had a longer AP duration in all measurements, AP rise time, AP decay time, and AP half-width (HW). AP rise time for single-firing hDRG-N was 1.045 ± 0.116ms and 1.265 ± 0.63ms for multi-firing hDRG-N, p=0.0143 (Figure 1R). AP decay time was also significantly longer in multi-firing hDRG-N at 10.64 ± 1.02ms and shorter in single-firing hDRG-N 4.934 ± 0.984ms, p<0.0001 (Figure 1S). AP HW was also longer in multi-firing hDRG-N at 4.483 ± 0.322ms and 2.545 ± 0.308ms, p<0.0001 (Figure 1T). We also looked at the maximum rise and decay slope. While there was no significant difference in max rise slope (Figure 1S), the max decay slope was significantly larger in multi-firing hDRG-N (-25.64 ± 1.61 dmV/dms) compared to single-firing hDRG-N (-43.14 ± 4.69 dmV/dms), p=0.0011 (Figure 1T).

### Correlations of electrophysiological properties in single-versus multi-firing hDRG-N

A Pearson correlation matrix was used to determine correlations of the electrophysiological properties measured in single- and multi-firing hDRG-N (Figure 2). Several properties were positively correlated with multi-firing cells including spontaneous activity when the cell was held at -45mV (SA_45), R_in_, FSL, visible sag, long AP duration (specifically decay time and HW), and a larger decay slope. Some properties were negatively correlated with multi-firing hDRG-N including cell capacitance and rheobase. We also found some interesting correlations throughout. For example, rheobase in hDRG-N was negatively correlated with AP rise, AP decay, and AP HW, suggesting that cells that have a long AP duration tend to fire a lower current input. AP rise time was negatively correlated with AP amplitude and AP peak showing that the length of the depolarization phase of an AP affects the amplitude of the AP. Capacitance was positively correlated with cell size which has been previously reported (Zheng et al., 2019). Taken together, these data help elucidate not only the electrophysiological properties of hDRG-N that are correlated with single and multi-firing cells, but also how these properties are correlated with other properties in hDRG-N.

**Figure 2.**
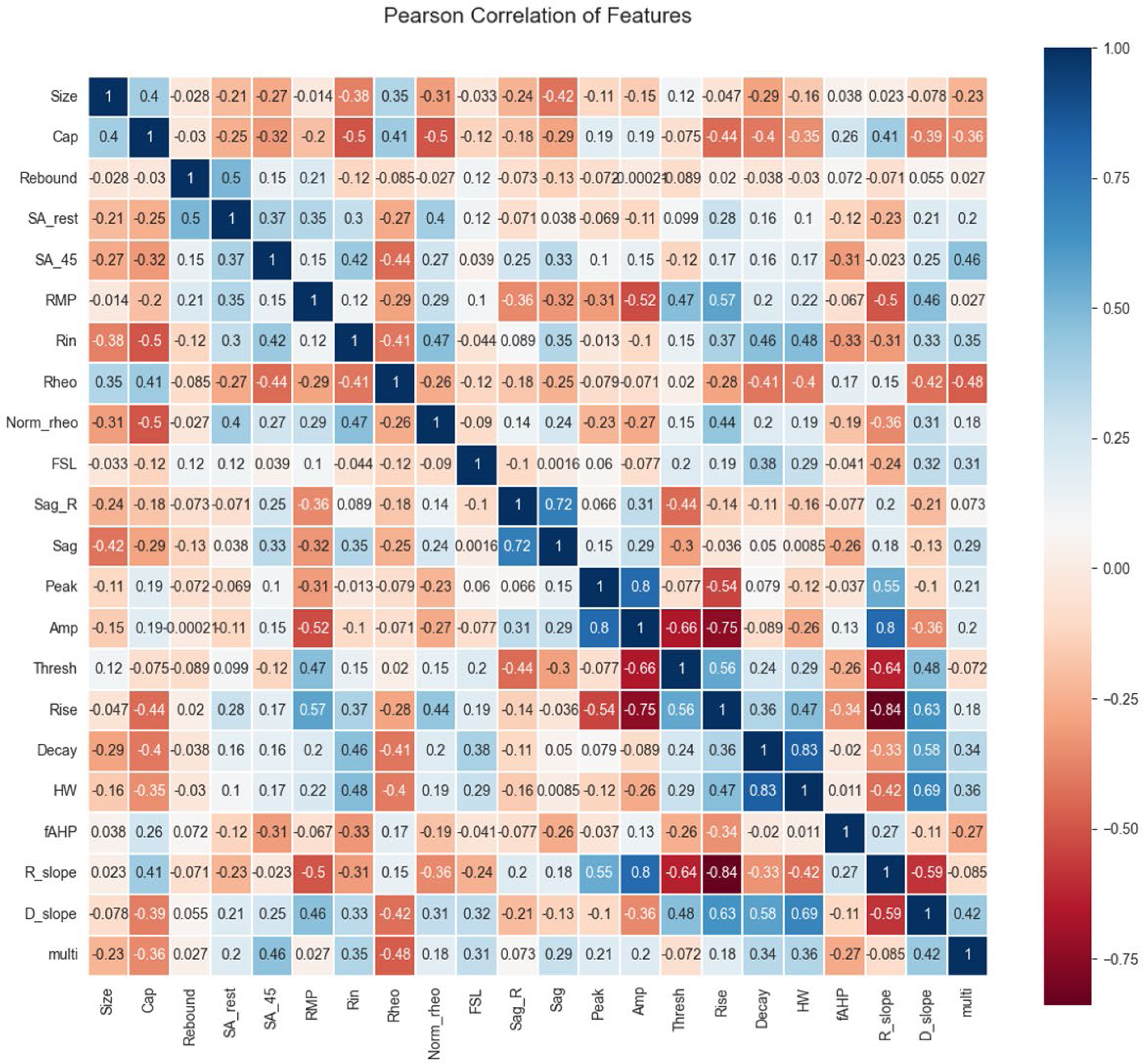
Pearson Correlation matrix of hDRG-N features. Features compared included size, capacitance (Cap), presence of rebound firing, spontaneous activity at rest (SA_rest), spontaneous activity at -45 mV (SA_45), resting membrane potential (RMP), input resistance (Rin), rheobase (Rheo), normalized rheobase (Norm_rheo), first spike latency (FSL), sag ratio (sag_R), presence of visible sag, AP peak, AP amplitude, AP threshold, AP rise time, AP decay, AP half-width, afterhyperpolarization (fAHP), rise slope (R_slope), decay slope (D_slope) and multi-firing.

### Comparison of model accuracies in predicting single- and multi-firing hDRG-N

To test if machine learning was able to predict multi-versus single firing cells based on electrophysiological features, we recorded from 94 hDRG-N from 10 donors. Since this is a relatively low number for machine learning models, we used a MCCV method that split the sample in an 80/20% train/test split across 1000 iterations. Then we ran all the splits through 5 different models, generated model performance scores, and extracted feature coefficients and importance’s from each model (Figure 3). We tested 5 machine learning models, logistic regression (LR), k-nearest neighbors (KNN), random forest classifier (RF), supported vector classification (SVC) and eXtreme Gradient Boost (XGBoost) with (Figure 4) and without feature selection (Figure S1, Figure S2). Machine learning models such as LR and SVC have a hard time handling highly correlated features, therefore we ran the models with all features (Figure S1, Figure S2) and with feature selection that accounts for highly correlated variables (Figure 4)(Ranganathan et al., 2017). Several features that are highly correlated are the features describing AP duration and AP amplitude. Since both AP rise and AP decay time effect AP HW and all three were significantly longer in multi-firing cells (Figure 1), we excluded AP rise time and AP decay time from the features used to determine model accuracy in our final models. Also, AP amplitude is highly correlated with AP threshold and AP peak. Since AP amplitude was not significantly different between single- and multi-firing cells but AP peak was, we decided to exclude AP amplitude and focus the models on AP peak. Rise slope and decay slope are direct measurements of the relationship of AP amplitude and AP duration, so those were also dropped from the final model predictions. Because our data set is relatively small for machine learning algorithms, we ran 1000 Monte Carlo simulations that randomized the 80/20 train/test split of the data. Model accuracies were plotted in a violin plot and model performance was compared (Figure 4A and B). We found that all models had accuracies around 80% on average (Figure 4B). SVC and XGBoost were the best performing models with and accuracy of 81.90% ± 0.29% and 81.88% ± 0.28% respectively. LR and KNN were the next best performing models with an accuracy of 80.30 ± 0.31% and 80.55 ± 0.31%. RF was the worst performing model with an accuracy of 76.91% ± 0.32%. RF is popular model in biological predictive algorithms, but our data shows that multiple models should be tested to determine the best model for a data set.

**Figure 3:**
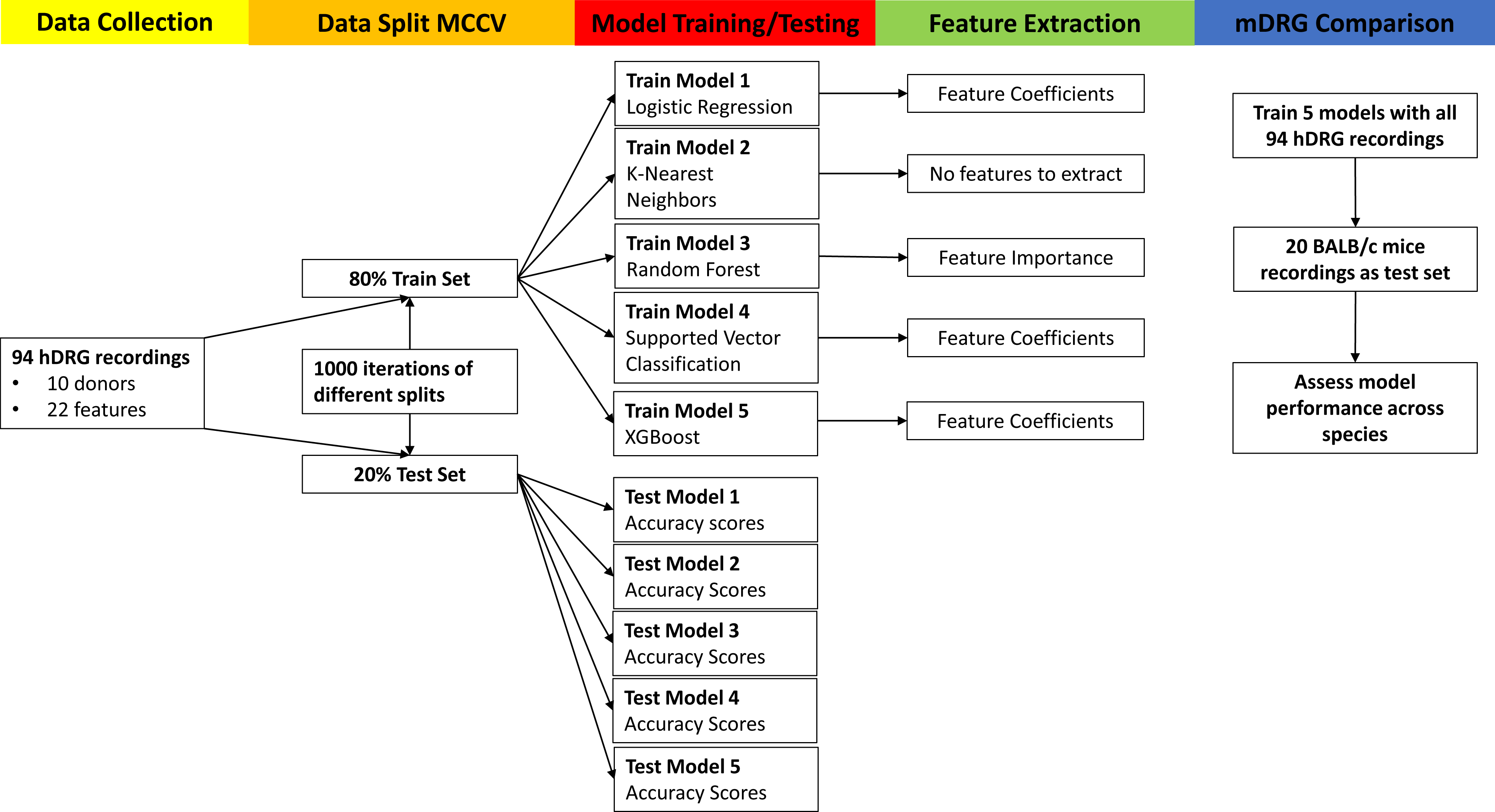
Machine Learning Model Flow Chart. Analysis was performed on 94 hDRG recordings, split using Monte Carlo Cross Validation (MCCV), 5 different models run, and features extracted. Finally, we ran this analysis again using recordings of mouse DRG (mDRG).

**Figure 4:**
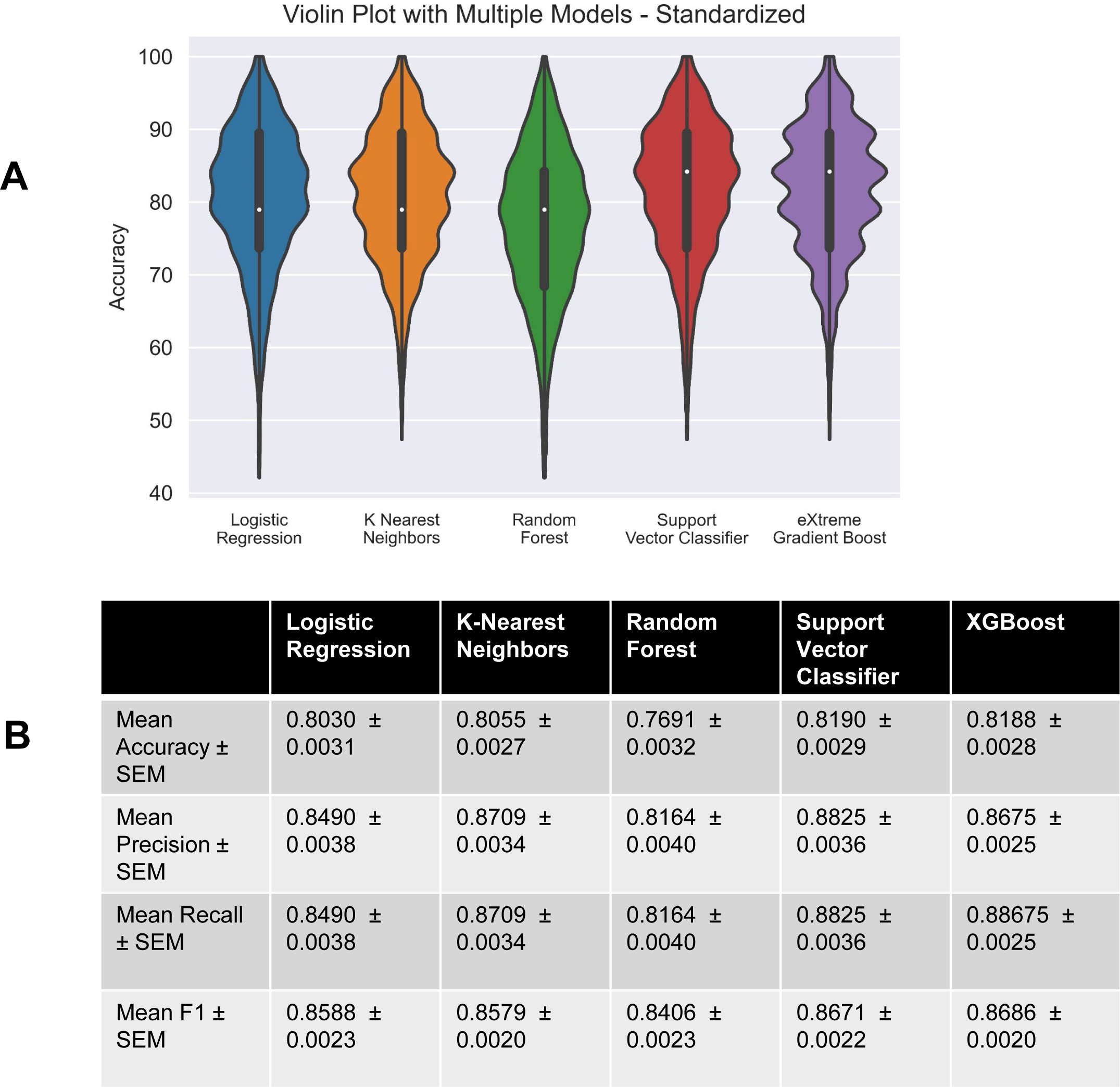
Performance of chosen models across randomized train-test splits. A. Accuracies of logistic regression, k-nearest neighbors, random forest, linear support vector machine, and eXtreme Gradient Boost on Monte Carlo Simulation (1000 randomly created 80-20 train-test splits) were compared with violin plots of model accuracy when trained and tested on standardized hDRG-N data. Plot represents the range of accuracies, and interior box and whisker plot shows quartiles and median. B. Table showing performance metrics of all models.

### Electrophysiological features that are predictive of single- and multi-firing hDRG-N

We next extracted feature coefficients for the LR and SVC models, and feature importance for the RF model (Figure 3). The KNN model is not able to compute to feature coefficients therefore it was not used to compute features predictive of multi-firing hDRG-N. Feature importance and coefficients are automatically calculated by the machine learning algorithms and can be extracted after model fitting. After running MCCV and generating our models, we extracted the resulting feature coefficients from the models to determine which features were most important in predicting single or multi-firing in hDRG-N. The LR, RF, and SVC model all predicted that a long FSL was the most important feature in predicting multi-firing cells while XGBoost predicted it as the third most important feature (Figure 5). XGBoost predicted a long AP duration (measured by HW) was the most important feature while LR and SVC predicted it as the second most important feature for predicting multi-firing hDRG-N, while the RF model predicted it as the third most important feature. Other features that had relatively high coefficients/importance in predicting multi-firing included rheobase, SA_45, AHP, and R_in_. We also used Shapley Additive Explanations to extract feature importance for all the models, and while the relative importance of each feature changed slightly, the same features, a long FSL, a long HW, SA_45, and a low rheobase all were scored highly in predicting multi-firing hDRG-N (Figure S3, Figure S4). Taken together, these data show that a long FSL and long HW are two of the most important features in predicting multi-firing hDRG-N.

**Figure 5:**
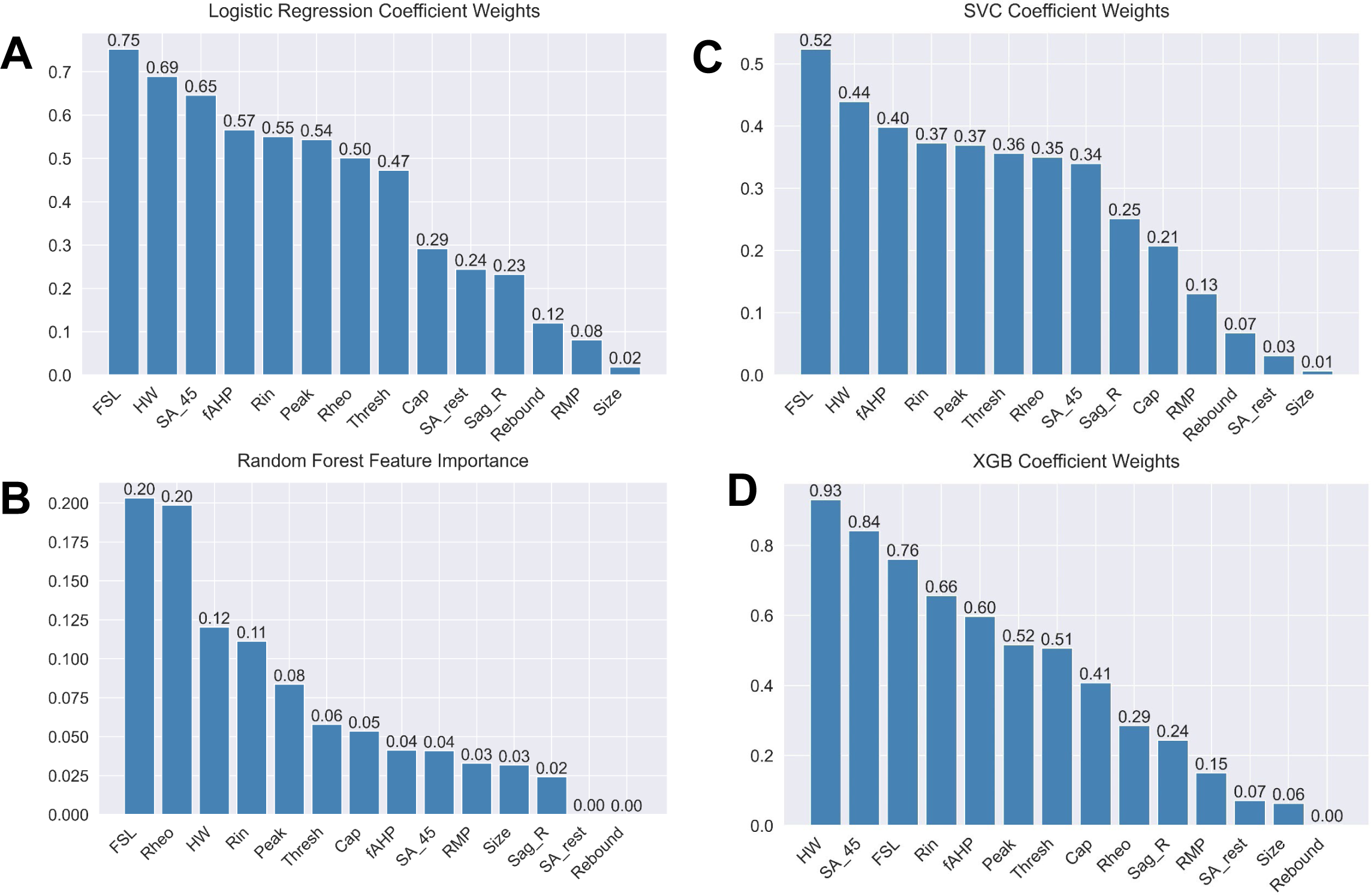
Feature importance metrics across logistic regression, random forest, and support vector classifier for standardized data. Mean coefficient weights or feature importance, when applicable, were extracted from the iterations in the Monte Carlo simulation and ordered. Absolute value of coefficients is displayed for logistic regression and SVC. A. Logistic regression B. Random forest C. Support vector classifier D. XGBoost

### Principal Component Analysis (PCA) extracts features important for multi-firing cells

We performed 3D PCA on our 94 hDRG-N using the features selected for the machine learning models. We found that there was a cluster of multi-firing cells that was distinct from the single-firing neurons (Figure 6A, orange dots), but some multi-firing cells did cluster with the single-firing cells (Figure 6A, blue dots). We found that 67.4% of the variance was explained by principal component 1(PC1), 23.8% of the variance was explained by PC2, and that 5.1% of the variance was explained by PC3 (Figure 6B). The feature importance contribution to each PC can be explained by the PC loadings. The loadings for PC1 were 0.73 for rheobase, 0.65 for R_in_, and 0.21 for FSL (Figure 6C). The feature loadings for PC2 were 0.88 for FSL, 0.43 for R_in_, and 0.12 for Rheo (Figure 6D). Since PC1 and PC2 explain 95% of the variance, the most important features for explaining the variance are rheobase, FSL, and R_in_.

**Figure 6:**
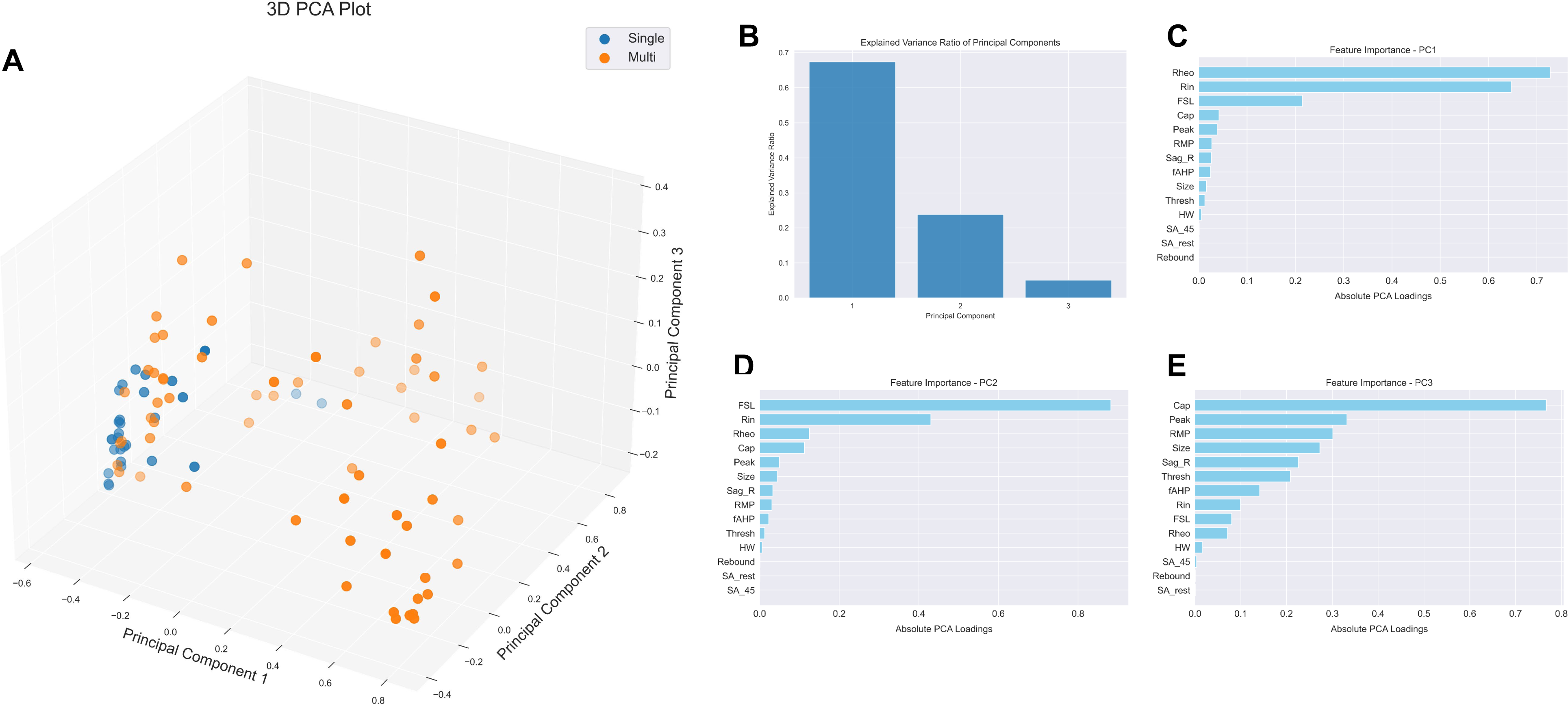
PCA with data normalization. A. PCA plot. B. Explained variance for each PC. C. Feature importance for PC1. D. Feature Importance for PC2. E. Feature importance for PC3.

### Machine learning models predictive of multi-firing in hDRG-N are able to predict multi-firing in mDRG-N

We found that similar to hDRG-N, ∼30% (6/20) of mDRG-N were single-firing in our data set. To test if the models and features that are important in single-versus multi-firing hDRG-N are also important in mDRG-N, we first compared the electrophysiological feature differences in single- and multi-firing cells in mDRG-N (Figure 7). We found several properties were significantly different in multi-firing mDRG-N such as a longer FSL, a lower rheobase, and a longer AP HW, similar to the properties that were different in hDRG-N. A Pearson Correlation Matrix shows that these features were strongly correlated with multi-firing cells (Figure 8). We found that the properties that were negatively correlated with multi-firing hDRG-N, such as rheobase, were also negatively correlated with mDRG-N, and that properties positively correlated with multi-firing hDRG-N, such as FSL and HW were also positively correlated in mDRG-N.

**Figure 7:**
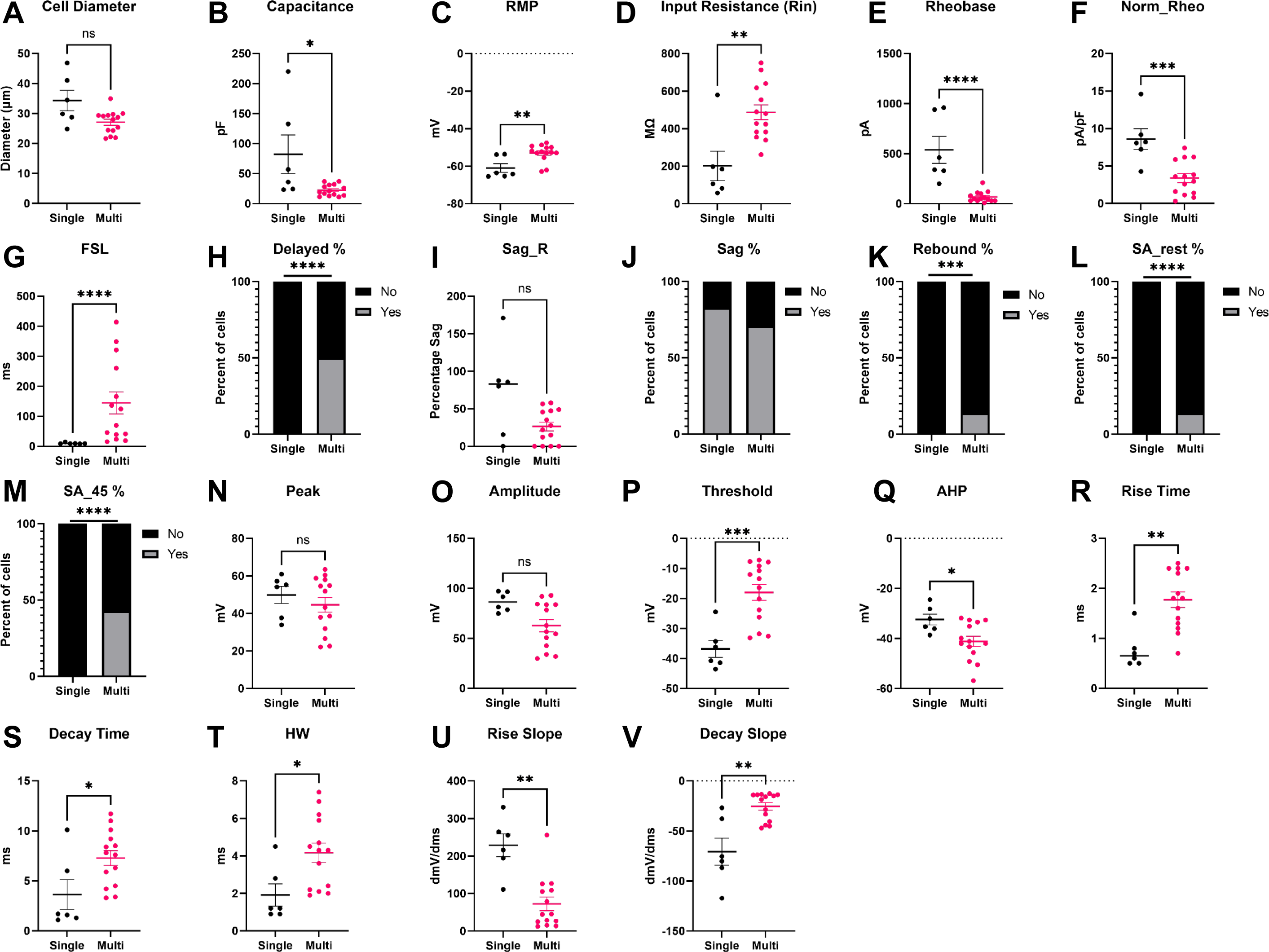
Intrinsic Properties and Firing Phenotypes of mDRG-N neurons in single versus multi-firing cells. Intrinsic properties and firing phenotypes were compared between single and multi-firing cells. A. Cell Diameter, B. Capacitance, C. Resting Membrane Potential (RMP), D. Input Resistance (Rin), E. Rheobase, F. Normalized Rheobase, G. First-Spike Latency (FSL), H. Percent delayed firing, FSL > 100ms, I. Sag Percentage at -100pA current input (Sag_R), J. Percentage cells with visible sag, K. Percentage of cells that have rebound firing. L. Percentage cells with spontaneous activity (SA) at rest, M. Percentage cells with spontaneous activity when held at -45mV. N. AP Peak. O. AP Amplitude. P AP Threshold. Q. AP After hyper-polarization (AHP). R. AP Rise Time, S. AP Decay Time, T. AP half-width. U. Max AP Rise Slope. V. Max AP Decay Slope. *p<0.05, **p<0.01, ***p<0.001, ****p<0.0001 by Mann-Whitney for A, B, C, D, E, F, G, I, N, O, P, Q, R, S, T, U, and V. By Fisher’s Exact test for H, J, K, L, and M .

**Figure 8.**
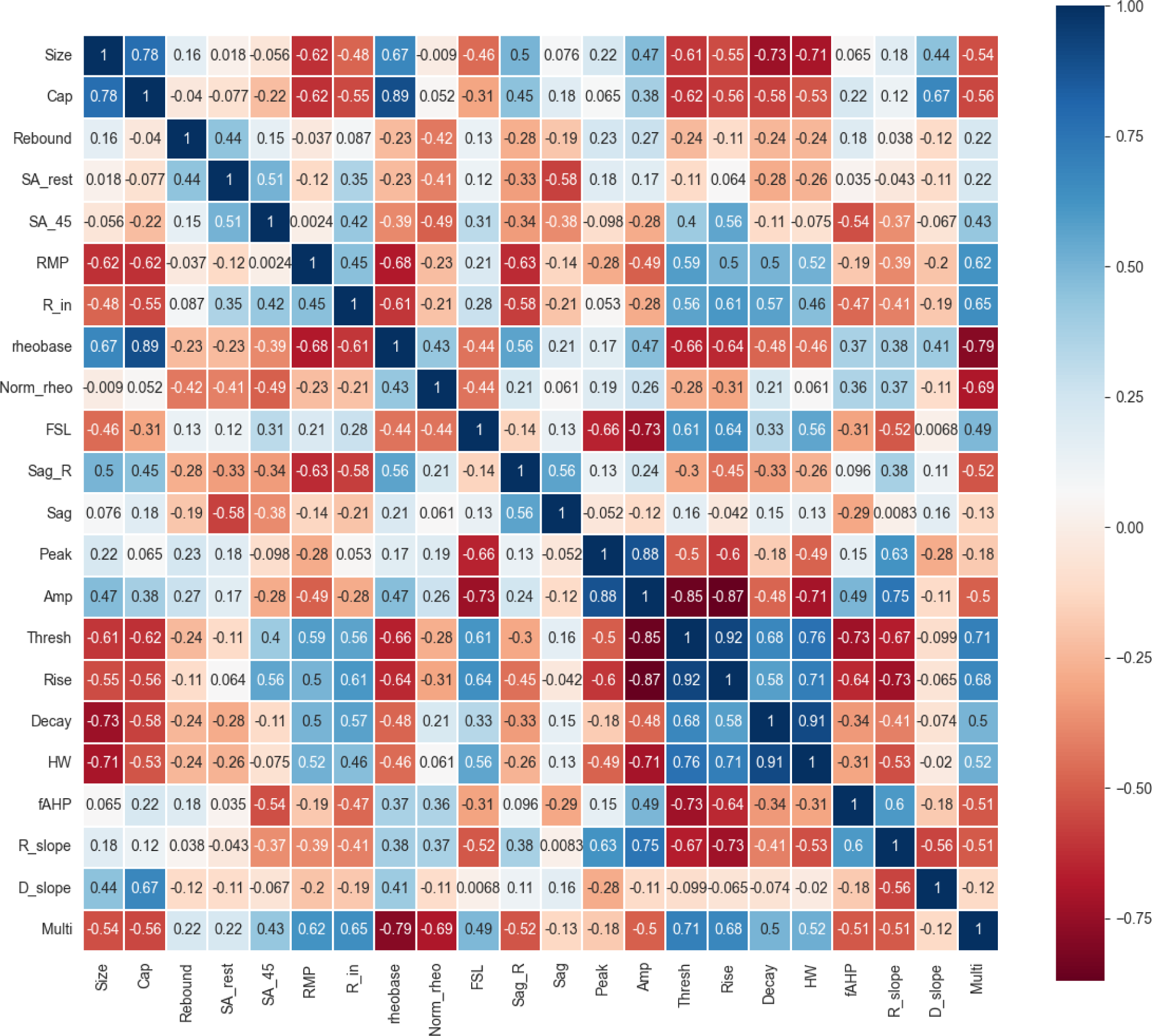
Pearson Correlation for mDRG properties. Features compared included size, capacitance (Cap), presence of rebound firing, spontaneous activity at rest (SA_rest), spontaneous activity at -45 mV (SA_45), resting membrane potential (RMP), input resistance (Rin), rheobase (Rheo), normalized rheobase (Norm_rheo), first spike latency (FSL), sag ratio (sag_R), presence of visible sag, AP peak, AP amplitude, AP threshold, AP rise time, AP decay, AP half-width, afterhyperpolarization (fAHP), rise slope (R_slope), decay slope (D_slope) and multi-firing.

Models generated using all 94 hDRG-N are able to predict single- and multi-firing in mDRG-N with greater than 90% Accuracy (Figure 9F). We found that the RF hDRG-N generated model performed the best when applied to the mDRG-N dataset with 100% accuracy, precision, recall, and F1 scores (Confusion Matrix, Figure 9C) while the SVC model performed second best with 95% accuracy, 93% precision, 100% recall, and 97% F1 score (Confusion Matrix, Figure 9D). LR, KNN, and XGBoost all performed similarly with 90% accuracy (Figure 9A, 9B, and 9F). These data show that the same electrophysiological features that predict multi-firing in hDRG-N are predictive of multi-firing in mDRG-N as well.

**Figure 9:**
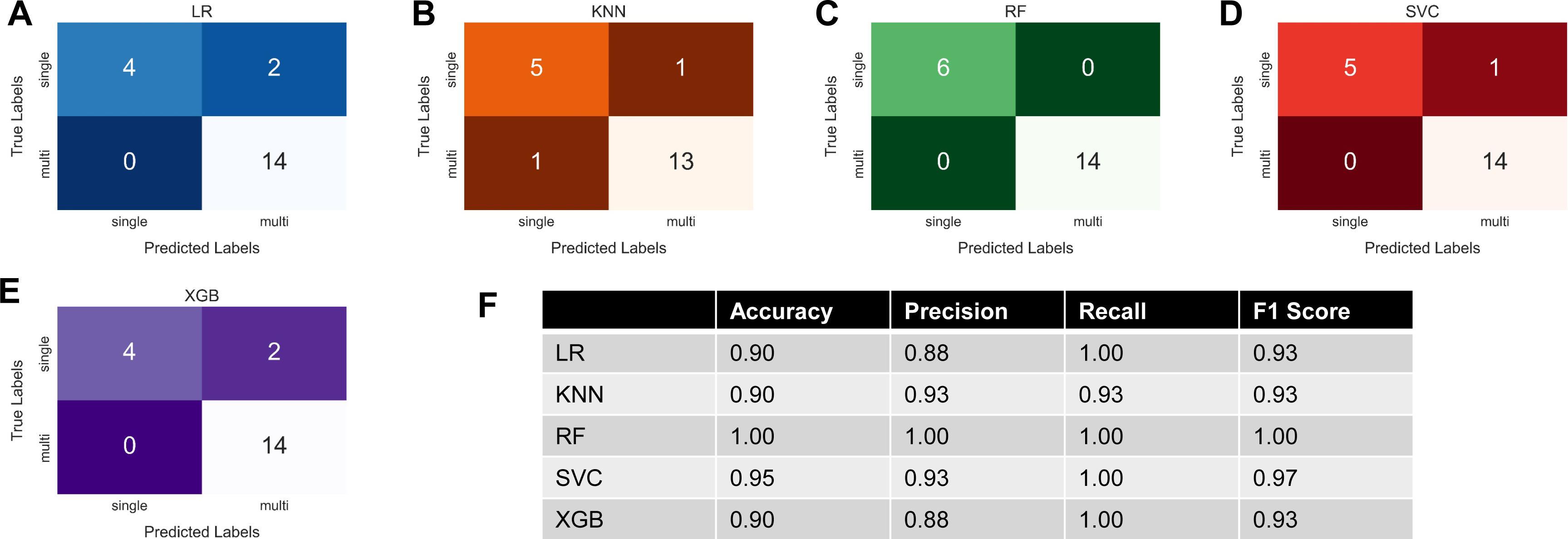
hDRG-N models are predictive of multi-firing in mDRG-N with high accuracy. 20 mDRG-N recordings were applied to models developed for hDRG-N. We are showing the A. Confusion matrix for LR model. B. Confusion matrix for KNN model. C. Confusion matrix for the RF model. D. Confusion matrix for SVC model. E. Confusion Matrix for XGBoost. F. Accuracy, precision, recall, and F-1 scores for mDRG-N compared to each model.

## Discussion

Both hDRG-N and mDRG-N are used to understand the electrophysiological properties associated with pain. Multi-firing DRG-N are more excitable than single-firing DRG-N; therefore, we examined the electrophysiological properties that are predictive of multi-firing using machine learning. We found that a similar percentage of neurons were multi-firing in both hDRG-N and mDRG-N, about 70%. We found that in both hDRG-N and mDRG-N, a long FSL and a long AP HW are the properties that are most predictive of multi-firing DRG-N. While many other features were significantly different in single-versus multi-firing cells, such as capacitance and AP peak, the machine learning shows that these specific features were not the most predictive of multi-firing. Machine learning helps us identify which features are likely to be important in regulating multi-firing in a way that standard statistical methods cannot. We developed our machine learning models using our hDRG-N data set which came from a diverse demographic of 10 donors. We found that several properties were negatively or positively correlated with multi-firing cells. For example, a long FSL, a long AP HW and a low rheobase were correlated with multi-firing cells. We also used a smaller dataset of naïve untreated BALB/c mice and found those same properties were correlated with multi-firing mDRG-N, though the correlations were much stronger. We hypothesize that since BALB/c share the same genetics and our sample looked only at naïve mice euthanized the same way, the properties of mDRG-N are more consistent when compared to our diverse set of hDRG-N with a variety of genetics, disease states, cause of death, and *in vivo* treatments. While there may be some differences in the molecular mechanisms governing multi- and single-firing in hDRG-N and mDRG-N, our data shows that the same electrophysiological properties are predictive of multi-firing across both species. In fact, the machine learning models we generated for the hDRG-N had higher accuracy scores when the mDRG-N data set was tested.

There are at least 8 neuronal subtypes in the DRG that can be defined by genetic labeling, nonpeptidergic nociceptors, peptidergic nociceptors, proprioceptors, and 5 classes of low threshold mechanoreceptors (LTMR), each with distinct firing patterns (Zheng et al., 2019). Several of these are multi-firing, such as nociceptors, C-LTMR, and proprioceptors, while some are single-firing, such as Aβ SA1-LTMR and Aβ Field-LTMR. The nociceptors and C-LTMR have very similar firing patterns, long AP HW and a long FSL, while the Aβ SA1-LTMR and Aβ Field-LTMR have a short AP HW and short FSL. Peptidergic DRG neurons that have a higher capacitance have a smaller AP HW and are less likely to be multi-firing (Zheng et al., 2019). While we only looked at single-firing versus multi-firing neurons, future work could focus on using machine learning to detect different neuronal subtypes based on firing patterns.

It has been reported by others that multi-firing hDRG-N have a longer FSL and longer AP HW (Yi et al., 2022). Molecular experiments in neurons have attributed changes in AP HW to a variety of voltage-gated sodium channels (Na_V_), voltage-gated potassium channels (K_v_), and voltage-gated calcium channels(VGCC), and that these ion channels are important in neuronal excitability and pain perception (Alles and Smith, 2018; Goodwin and McMahon, 2021). Several Na_V_ channels have been implicated in pain in humans and animal models, including Na_V_1.7 and Na_V_1.8 (Alles and Smith, 2021; Goodwin and McMahon, 2021). Na_V_1.7 (HWTX-IV) and Na_V_1.8 (A-803467, PF-01247324) antagonists decrease neuronal excitability of sensory neurons (Payne et al., 2015; Ye et al., 2015; Atmaramani et al., 2020; Mulpuri et al., 2022), while activation of Na_V_1.8 increases neuronal excitability (Ye et al., 2015). Sensory neurons with increased Na_V_1.8 expression had longer AP HW (Djouhri et al., 2003; Thériault and Chahine, 2014; Zheng et al., 2019; Mulpuri et al., 2022). Knockdown of the β4 subunit of Na_V_1.8 decreased AP HW and decreased neuronal excitability in mDRG-N (Xiao et al., 2019). Single cell RNA-seq has found that increased Na_V_1.8 and Na_V_1.9 expression, and decreased Na_V_1.7 expression are correlated with an increase in AP HW (Thériault and Chahine, 2014). Expression of Na_V_1.8 and Na_V_1.9 is high in multi-firing nociceptors and C-LTMR, and low in single-firing Aβ-LTMR (Zheng et al., 2019). Taken together these data suggest that Na_V_1.8, and possibly Na_V_1.9 increase AP HW and neuronal excitability, and that targeting Na_V_1.8 pharmacologically decreases AP HW and excitability. Our machine learning models show that increased AP HW is predictive of multi-firing cells and the literature suggests Na_V_1.8 may be a molecular mechanism that regulates both AP HW and neuronal excitability.

Several K_v_ channels are also implicated in regulating AP HW, FSL, and neuronal excitability. Blocking K_v_3 And K_v_1 channels with TEA and DTX increased the AP HW in hippocampal neurons (Hoppa et al., 2014). Experiments in rat DRG neurons show that dominant-negative mutation in K_v_3.4 effectively broadens AP HW showing that K_v_3.4 governs AP repolarization (Alexander et al., 2022). Rat DRG-N treated with calcineurin which attenuates (decreases) Kv3.4 action, increases AP HW (Zemel et al., 2017). Data shows that multi-firing nociceptors and C-LTMR have a long AP HW and a long FSL (Zheng et al., 2019). These neuronal subtypes showed decreased levels of K_v_1.1, K_v_1.2, K_v_2.1, K_v_3.1, K_v_3.3, K_v_7.2, K_v_7.3, K_v_9.1, and K_v_11.1 and increased levels of K_v_4.1, K_v_6.2 and K_v_6.3 when compared to single firing Aβ-LTMR (Zheng et al., 2019). Using a variety of K_v_ inhibitors, the main contribution to potassium currents in nonpeptidergic nociceptors were K_v_1, K_v_2, and K_v_4, while in C-LTMR the main contributor was K_v_4 (Zheng et al., 2019). In single-firing Aβ-LTMR the main contributor to potassium currents was K_v_3 followed by K_v_1 (Zheng et al., 2019). Computational modeling showed that C-LTMR neurons without Kv4 would have a short FSL. Blocking K_v_4.3 with AmmTx3 or K_v_4.3 KO mice had a decreased FSL in C-LTMR, while blocking K_v_1 with DTx had no effect on firing properties in C-LTMR (Zheng et al., 2019). Blocking K_v_1 current with DTx in single firing Aβ-LTMR turned them into multi-firing neurons (Zheng et al., 2019).

Calcium channels may also be involved in regulating FSL, AP HW, and neuronal excitability. The voltage-gated calcium channel alpha-2-delta-1 subunit (α_2_δ-1) has been implicated in chronic pain and neuronal excitability (Newton et al., 2001; Alles and Smith, 2018; Dolphin, 2018; Cui et al., 2021). Experiments in mDRG-N show that DRG cultured from α_2_δ-1 knock out mice have a shorter FSL and AP HW and that these neurons are less excitable, i.e. firing fewer APs (Margas et al., 2016). Ca_V_2.2 and α_2_δ-1 expression is also increased in multi-firing nociceptors and C-LTMR with long AP HW and long FSL, while it is absent in single Aβ-LTMR with short AP HW and short FSL (Zheng et al., 2019). Gabapentinoids are used as first line treatment for neuropathic pain and act on α_2_δ-1 (Alles and Smith, 2018). Many publications show treatment with gabapentinoids decrease evoked calcium currents and neuronal excitability (Bannister et al., 2011; Biggs et al., 2014, 2015). The calcitonin gene related peptide (CGRP) was also increased in multi-firing nociceptors and C-LTMR (Zheng et al., 2019).

There are a lot of data that suggest a variety of ion channels affecting AP HW and FSL. Our machine learning suggests that these are the most predictive features of single-versus multi-firing in DRG-N. Much of the work to elucidate which ion channels are important for these features has been done in rodents and not hDRG-N. Given the machine learning models for hDRG-N are also able to predict multi-firing in mDRG-N, and that the same electrophysiological features are correlated with multi-firing cells in both species, we predict the same ion channels would be important in hDRG-N. To test this hypothesis, single cell patch-RNA-seq could be used to determine which ion channels are up- or down-regulated in multi-firing hDRG-N. Additionally, it has been shown that there are at least 8 neuronal subtypes in DRG-N, with a variety of firing patterns that can be differentiated by genetic markers. Our current dataset and machine learning tools cannot differentiate these neuronal subtypes presently. However, we hope to continue expanding our capabilities in future studies. Single-cell RNA-seq and post-hoc labeling could be used to differentiate these neuronal subtypes in hDRG-N and future machine learning models could be applied to these data.

## Supplementary Figure Legends

**Figure S1.**
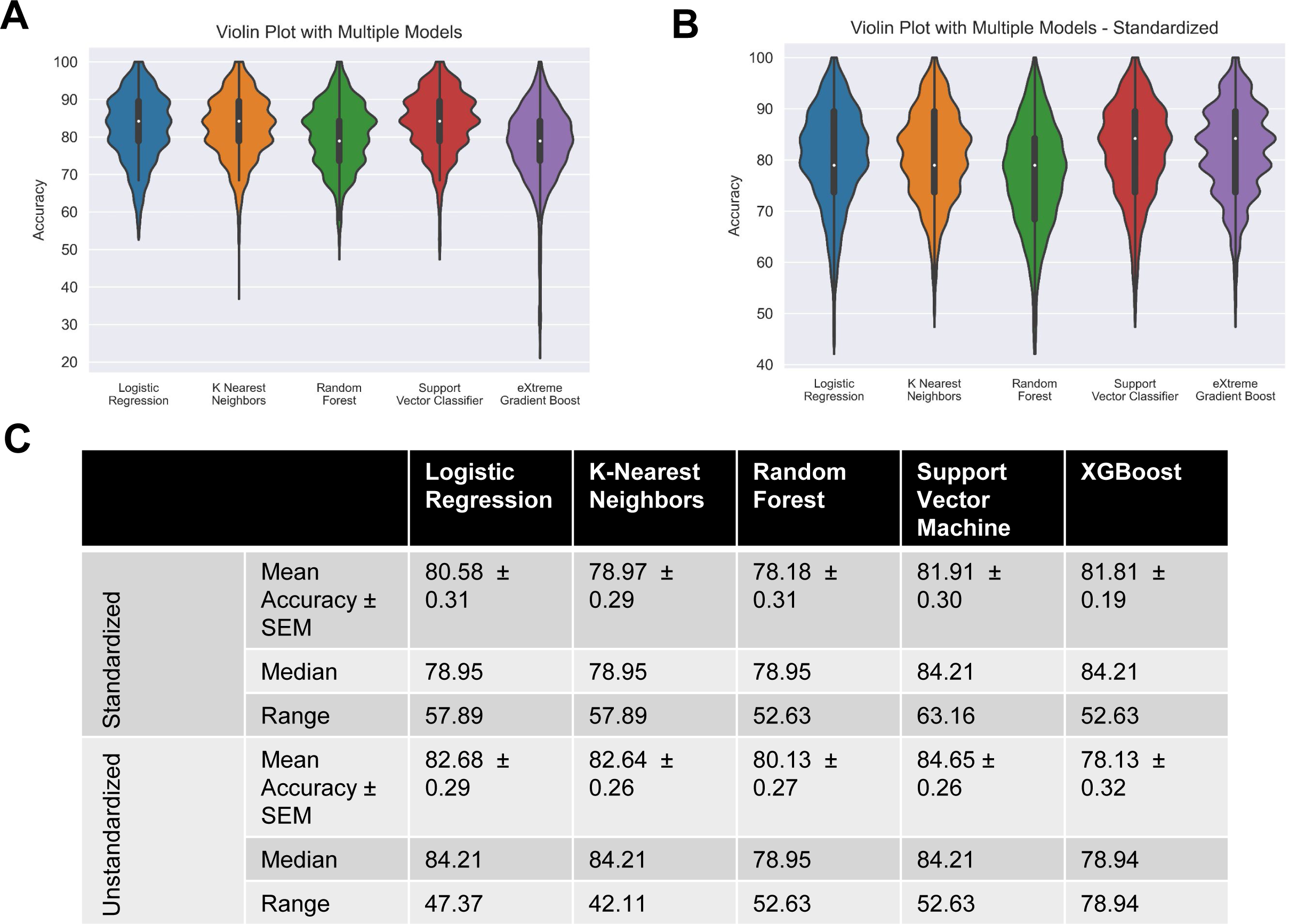
Performance of chosen models across randomized train-test splits. Performances of logistic regression, k-nearest neighbors, random forest, and linear support vector machine on Monte Carlo Simulation (1000 randomly created 80-20 train-test splits) were compared with violin plots of model accuracy when trained and tested on standardized/unstandardized dataset, hue represents model, plot is cut to represent the range of accuracies, and interior box and whisker plot shows quartiles and median. A. Unstandardized dataset B. Standardized dataset C. Table showing performance metrics of both groups of models

**Figure S2:**
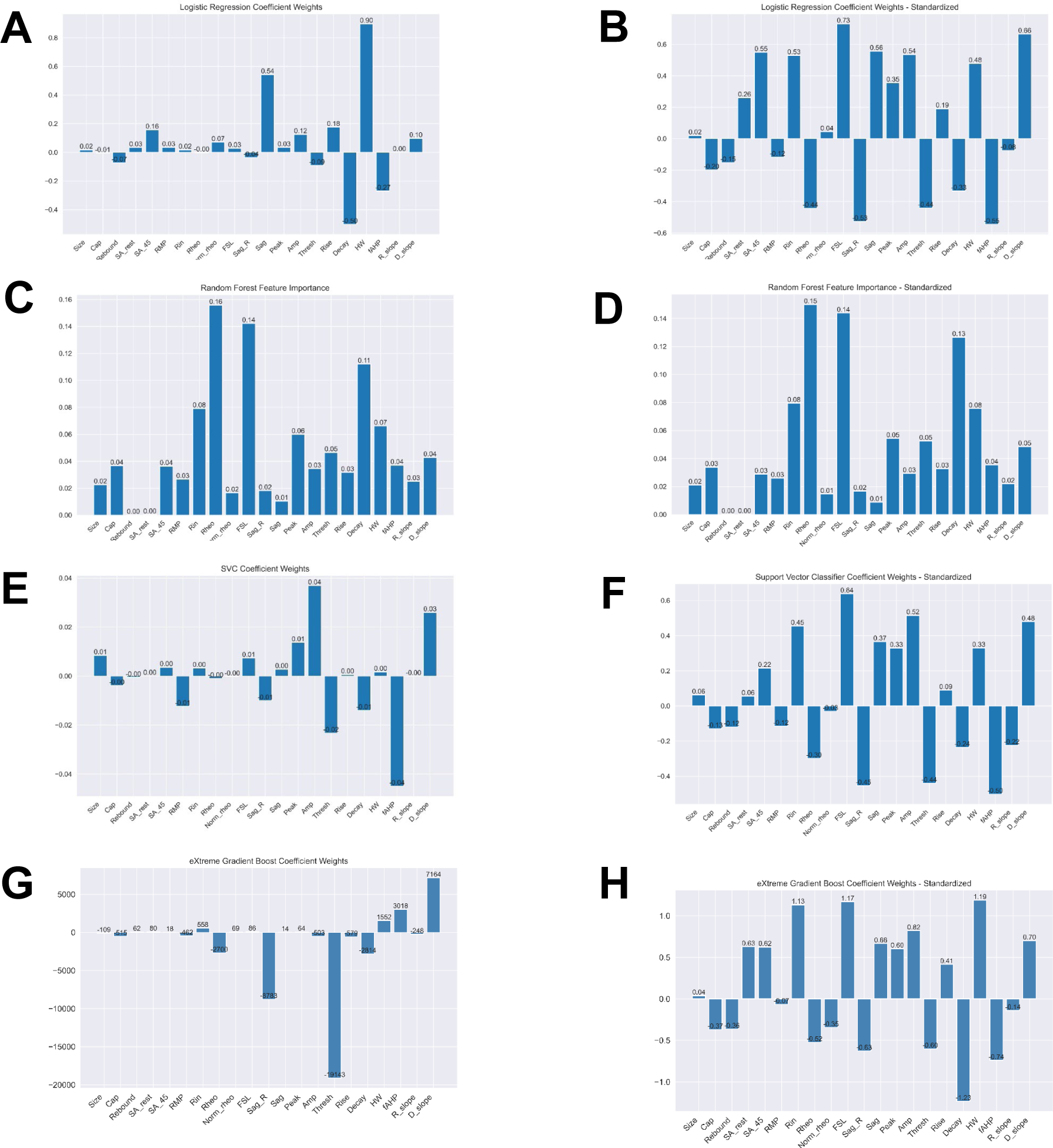
Feature importance metrics across logistic regression, random forest, and support vector classifier for standardized and unstandardized data. Mean coefficient weights or feature importance, when applicable, were extracted from the iterations in the Monte Carlo simulation. Graphs on left represent models with unstandardized data while right use standardized A-B. Logistic regression C-D. Random forest E-F. Support vector classifier

**Figure S3:**
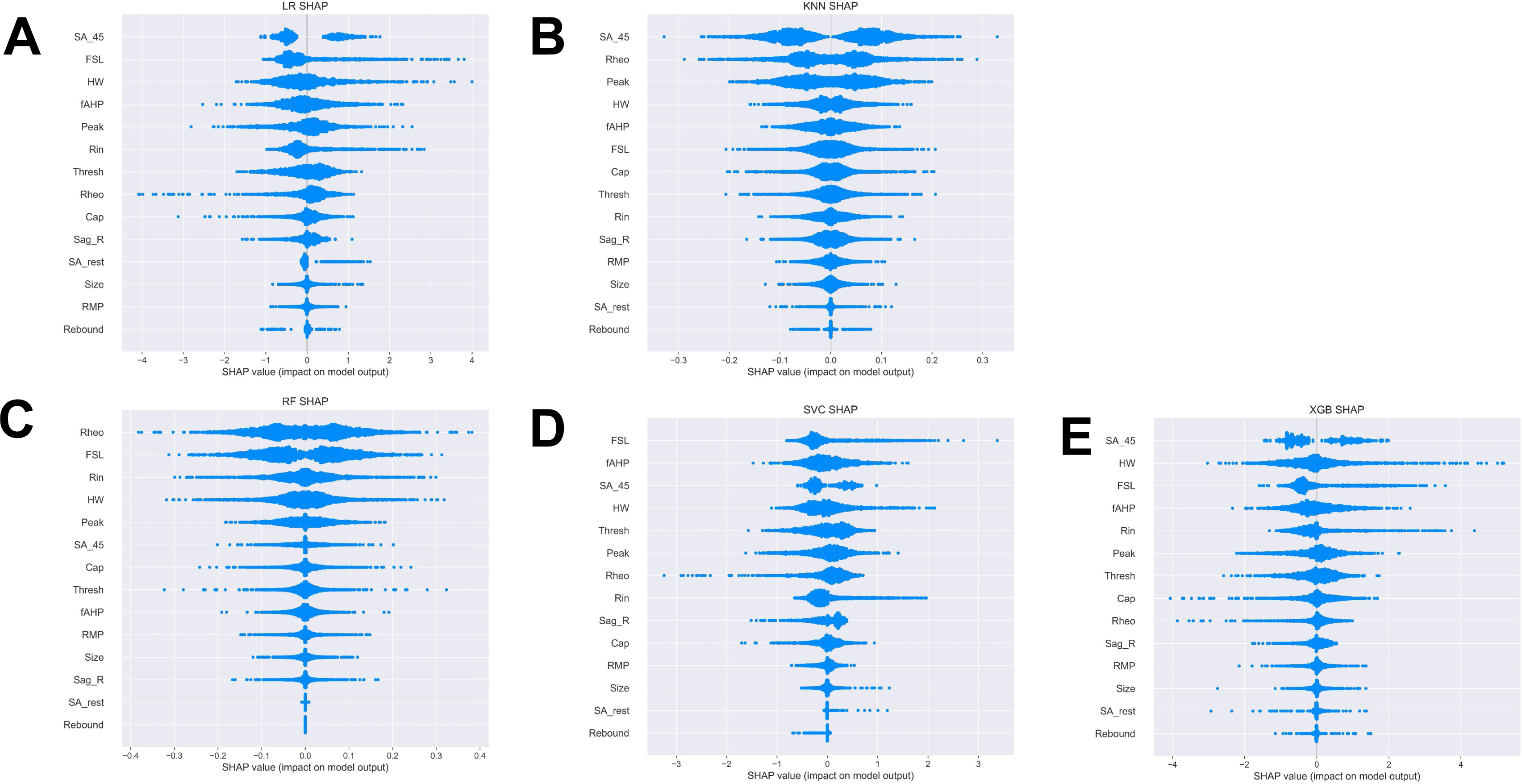
Monte Carlo Shapely Additive Explanations (SHAP) Beeswarm Plots. Beeswarm plots for A. Logistic Regression. B. K-Nearest Neighbors. C. Random Forest. D. Supported Vector Classification. E. XGBoost.

**Figure S4:**
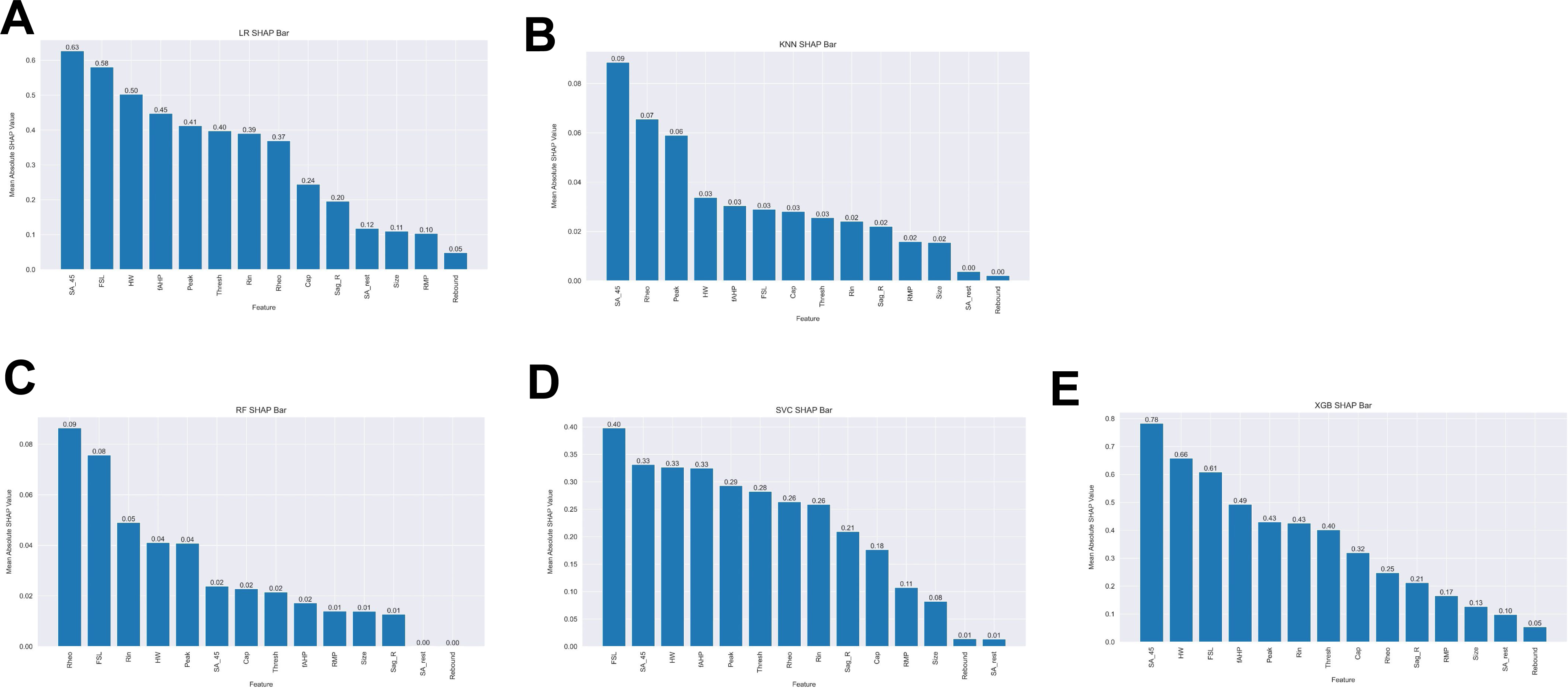
Monte Carlo Shapely Additive Explanations (SHAP) Values. A. Logistic Regression. B. K-Nearest Neighbors. C. Random Forest. D. Supported Vector Classification. E. XGBoost.

## Disclosures

This study was supported by NIH 1UG3NS123958-01 (K.N.W., S.R.A.A.), associated NIH Diversity Supplement 3UG3NS123958-01S1 (A.E.G.), and the Research Endowment Fund of the Department of Anesthesiology and Critical Care Medicine, University of New Mexico Health Sciences Center.

## Acknowledgements

The authors are grateful to the donors and their families and humbled to have the opportunity to perform this research. We thank the organ donation and transplant team, who have supported and facilitated this project in numerous ways.

## Notes

### Competing Interest Statement

The authors have declared no competing interest.

